# Formation of secondary allo-bile acids by novel enzymes from gut Firmicutes

**DOI:** 10.1101/2022.08.09.503364

**Authors:** Jae Won Lee, Elise S. Cowley, Patricia G. Wolf, Heidi L. Doden, Tsuyoshi Murai, Kelly Yovani Olivos Caicedo, Lindsey K. Ly, Furong Sun, Hajime Takei, Hiroshi Nittono, Steven L. Daniel, Isaac Cann, H. Rex Gaskins, Karthik Anantharaman, João M. P. Alves, Jason M. Ridlon

## Abstract

The gut microbiome of vertebrates is capable of numerous biotransformations of bile acids, which are responsible for intestinal lipid digestion and function as key nutrient-signaling molecules. The human liver produces bile acids from cholesterol predominantly in the A/B-*trans* orientation in which the sterol rings are “kinked”, as well as small quantities of A/B-*cis* oriented “flat” stereoisomers known as “primary allo-bile acids”. While the complex multi-step bile acid 7α-dehydroxylation pathway has been well-studied for conversion of “kinked” primary bile acids such as cholic acid (CA) and chenodeoxycholic acid (CDCA) to deoxycholic acid (DCA) and lithocholic acid (LCA), respectively, the enzymatic basis for the formation of “flat” stereoisomers allo-deoxycholic acid (allo-DCA) and allo-lithocholic acid (allo-LCA) by Firmicutes has remained unsolved for three decades. Here, we present a novel mechanism by which Firmicutes generate the “flat” bile acids allo-DCA and allo-LCA. The BaiA1 was shown to catalyze the final reduction from 3-oxo-allo-DCA to allo-DCA and 3-oxo-allo-LCA to allo-LCA. Phylogenetic and metagenomic analyses of human stool samples indicate that BaiP and BaiJ are encoded only in Firmicutes and differ from membrane-associated bile acid 5α-reductases recently reported in Bacteroidetes that indirectly generate allo-LCA from 3-oxo-Δ^4^-LCA. We further map the distribution of *baiP* and *baiJ* among Firmicutes in human metagenomes, demonstrating an increased abundance of the two genes in colorectal cancer (CRC) patients relative to healthy individuals.

**SIGNIFICANCE STATEMENT:** Bile acid synthesis by vertebrates is central to digestion and nutrient signaling. Gut bacteria have evolved enzymes capable of converting primary bile acids to hundreds of secondary bile acids. While bile acid microbiology has been focused on the metabolism of ring hydroxyl groups and the carboxylated side-chain, very little is known about how bacteria alter the shape of the steroid ring system. Here, we describe enzymes expressed by Firmicutes that convert the “kinked” primary bile acid into “flat” secondary bile acids. Decades of research indicate that increased levels of secondary bile acids are risk factors for colorectal cancer. Hidden Markov Models developed from the BaiP and BaiJ enzyme sequences revealed significant enrichment in metagenomes of subjects with colorectal cancer.

## INTRODUCTION

Bile acid synthesis in the liver and represents a major route for removal of cholesterol from the body and bile acids function as an emulsifying agent for the digestion of lipid-soluble dietary components in the aqueous lumen of the small bowel (1). In humans, the liver synthesizes two abundant primary bile acids, cholic acid (CA; 3ɑ-,7ɑ-,12ɑ-trihydroxy-5β-cholan-24-oic acid) and chenodeoxycholic acid (CDCA; 3ɑ-,7ɑ-dihydroxy-5β-cholan-24-oic acid) from cholesterol. Before active secretion from the liver, bile acids are conjugated to either taurine or glycine at the C-24 carboxyl group (1). When bile acids reach the terminal ileum, they are actively transported across the epithelium into portal blood and returned to the liver in a process known as enterohepatic circulation (EHC). Several hundred milligrams of bile acids escape EHC daily and enter the large intestine. Colonic bacteria are capable of carrying out numerous biotransformations of primary bile acids to diverse secondary bile acids in the large intestine. The composition of intestinal and fecal bile acids in germ-free animals reflects the biliary composition (2-5). Meanwhile, in conventional animals with a normal gut microbiota, fecal bile acids composition is diversified from only a few primary bile acids derived from the host to an estimated ∼400 secondary bile acid products (6, 7). Bacterial modifications to bile acids provides a form of interdomain communication given that beyond mere lipid-digesting detergents, bile acids are important nutrient-signaling molecules (8). Indeed, microbial metabolism of bile acids is widely recognized to contribute to numerous human disorders including, but not limited to, cancers of the liver (9, 10) and colon (11), obesity, type 2 diabetes, non-alcoholic fatty liver disease (NAFLD) (12, 13), cholesterol gallstone disease (14, 15), Alzheimer’s disease (16, 17), and cardiovascular disease (18).

A myriad of microbial bile acid biotransformations occur in the large intestine and include two key transformations. First, the conjugated bile acids are hydrolyzed to unconjugated bile acids and glycine or taurine by bile salt hydrolase (BSH). Second, the unconjugated primary bile acids CA and CDCA are converted to deoxycholic acid (DCA; 3ɑ-,12ɑ-dihydroxy-5β-cholan-24-oic acid) and lithocholic acid (LCA; 3ɑ-hydroxy-5β-cholan-24-oic acid) (19) via 7ɑ-dehydroxylation, respectively. BSH (EC 3.5.1.24) enzymes are widely distributed among predominant microbial phyla within the domains Bacteria and Archaea inhabiting the human GI tract and catalyze the substrate-limiting deconjugation of bile acid amides (20). The resulting major secondary bile acids routinely measured in human fecal samples are unconjugated derivatives of DCA and LCA (20). A bile acid inducible (*bai*) regulon encoding *bai* enzymes involved in the conversion of CA to DCA (**Fig. 1**), and CDCA and ursodeoxycholic acid (UDCA; 3ɑ-,7β-dihydroxy-5β-cholan-24-oic acid) to LCA has been elucidated over the past three decades in strains of *Lachnoclostridum scindens* (formerly *Clostridium scindens*), *Peptacetobacter hiranonis* (formerly *Clostridium hiranonis*), and *Lachnoclostridum hylemonae* (formerly *Clostridium hylemonae*) (21). Discovery and characterization of *bai* genes has allowed recent studies to extend the species distribution of 7-dehydroxylating bacteria into new families within the Firmicutes through bioinformatics-based searches of metagenomic sequence databases (22, 23). Similarly, comparison of the distribution of *bai* genes between fecal metagenomes obtained from healthy and disease cohorts has also enabled the association of the abundance of *bai* genes with risk for adenomatous polyps (24) or colorectal cancer (25). This agrees with bile acid metabolomic studies that demonstrate increased fecal and serum DCA and LCA derivatives in subjects at high risk for CRC (26-31). Conversely, lower abundance of *bai* genes is associated with bile acid dysbiosis characterized by increased fecal conjugated primary bile acids in inflammatory bowel diseases (32, 33).

**Fig. 1.**
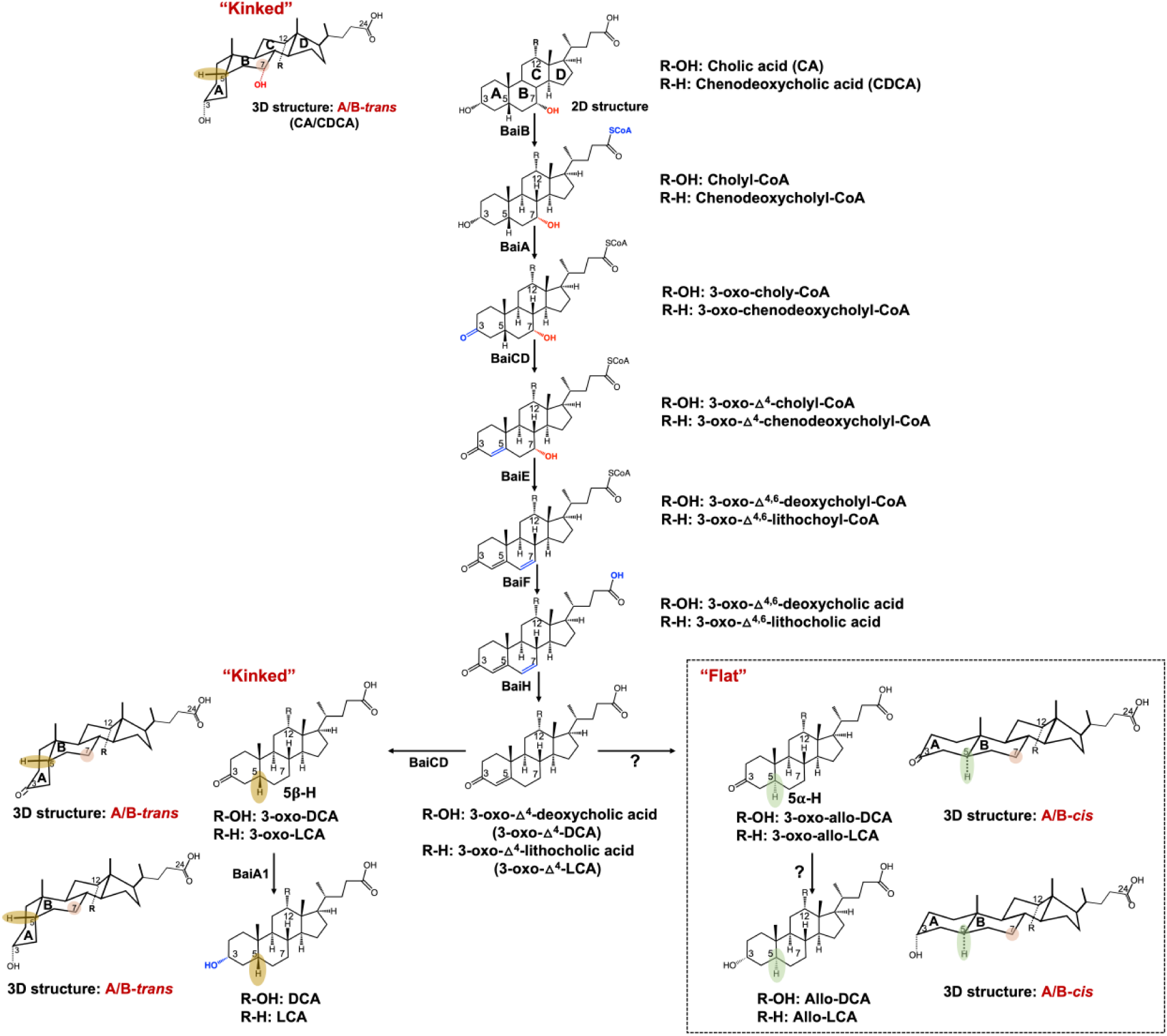
A proposed pathway for the 7α-dehydroxylation of cholic acid (CA) and chenodeoxycholic acid (CDCA) to deoxycholic acid (DCA) and allo-deoxycholic acid (allo-DCA), and lithocholic acid (LCA) and allo-lithocholic acid (allo-LCA). BaiB, Bile acid CoA ligase; BaiA, 3α-hydroxysteroid dehydrogenase; BaiCD, 3-dehydro-Δ^4^-7α-oxidoreductase; BaiE, 7α-dehydratase; BaiF, CoA transferase; BaiH, 3-dehydro-Δ^4^-7β-oxidoreductase. The enzymes involved in the sequential reduction of 3-oxo-4-DCA to 3-oxo-allo-DCA and allo-DCA are currently unknown.

There are additional *bai* genes yet to be accounted for in strains of *L. scindens* that result in the formation of stereoisomers of DCA and LCA known as “secondary allo-bile acids”. In 1991, Hylemon et al. (34) reported that allo-deoxycholic acid (allo-DCA; 3ɑ-,12ɑ-dihydroxy-5ɑ-cholen-24-oic acid) formation is a CA-inducible side-product of bile acid 7-dehydroxylation by *L. scindens*. During the conversion of cholesterol to the primary bile acids CA and CDCA, the liver enzyme Δ^4^-3-ketosteroid-5β-reductase (3-oxo-Δ^4^-steroid-5β-reductase; AKR1D1) saturates the Δ^4^-bond generating steroid A/B rings in the *cis-*orientation which appear “kinked” (**Fig. 1**). When CA is transported into bacteria expressing *bai* genes, the first oxidative steps of bile 7-dehydroxylation, catalyzed by BaiA and BaiCD, “resetting” A/B ring stereochemistry through formation of the 3-keto-Δ^4^ structure. This is followed by the rate-limiting 7ɑ-dehydration (BaiE) (35). The BaiCD was shown to then re-establish stereochemistry by catalyzing the conversion of 3-oxo-Δ^4^-DCA (12ɑ-hydroxy-3-oxo-5β-chol-4-en-24-oic acid) to 3-oxo-DCA (12ɑ-hydroxy-3-oxo-5β-cholan-24-oic acid), which is further reduced by BaiA1 and BaiA2 to DCA. The current model of bile acid 7ɑ-dehydroxylation suggests that another enzyme, currently unknown, acts on 3-oxo-Δ^4^-DCA to form the alternative stereoisomer, 3-oxo-allo-DCA (12ɑ-hydroxy-3-oxo-5ɑ-cholan-24-oic acid), which is reduced by another unknown reductase to allo-DCA. Secondary allo-bile acids have a “flat” shape owing to hydrogenation that results in an A/B-*trans* orientation (**Fig. 1**). While few studies have reported measurement of allo-DCA and allo-LCA (3-oxo-5ɑ-cholan-24-oic acid), two studies have shown these bile acids are enriched in the feces of patients with CRC (36, 37). Derivatives of allo-LCA are also reported to be enriched in Japanese centenarians (38), although there is a paucity of measurement of secondary allo-bile acids across populations and disease states. Thus, determining the gene(s) encoding reductases in *L. scindens* and other gut microbes responsible for the formation of allo-DCA and allo-LCA is of biomedical importance.

We recently reported genome-wide transcriptome profiling of *L. scindens* ATCC 35704 in the presence of CA and DCA and identified a potential candidate bile acid-inducible 3-oxo-Δ^4^-5ɑ-reductase (39). Here, we confirm that this candidate bile acid-inducible gene encodes a novel bile acid 3-oxo-Δ^4^-5ɑ-reductase responsible for secondary allo-bile acids formation. We have named this gene in *L. scindens* ATCC 35704 the *baiP* gene. We previously reported identification of the *baiJ* gene as part of a polycistronic operon in *L. scindens* VPI 12708 and *L. hylemonae* TN271, whose function remained unknown (40). Our current study reports that the *baiJ* gene also encodes a bile acid 3-oxo-Δ^4^-5ɑ-reductase. The *baiP* and *baiJ* genes are distributed solely among the Firmicutes. Identification of these *bai* genes may provide the ability to predict and potentiate the formation of alternative forms of secondary bile acids whose ring structures are “flat” rather than the “kinked” form produced by the host. Indeed, we developed Hidden Markov Models (HMMs) of *bai* proteins and determined the distribution of *baiP* and *baiJ* in human metagenomes, demonstrating increased abundance in colorectal cancer (CRC) patients relative to healthy individuals.

## RESULTS

### The HDCHBGLK_03451 gene from *L. scindens* ATCC 35704 encodes a bile acid 5ɑ-reductase, yielding secondary allo-bile acids

Prior work established that allo-DCA is a CA-induced side-product of CA metabolism in cell-extracts of *L. scindens* VPI 12708 (34) (**Fig. 1**). We previously identified *L. scindens* ATCC 35704 gene HDCHBGLK_03451 as CA-inducible and suggested this is a likely candidate for bile acid 5ɑ-reductase (39) (**Fig. 2a**). The gene HDCHBGLK_03451 encodes a 563 amino acid protein comprising FMN (flavin mononucleotide) and FAD (flavin adenine dinucleotide)-binding domains (**Fig. 2b**). The HDCHBGLK_03451 gene from *L. scindens* ATCC 35704 was codon-optimized for *E. coli* and overexpressed in *E. coli* (**Fig. 2c**) for resting cell assays with bile acid intermediates (**Fig. 2d**). The stereochemistry of the A/B ring junction is lost during the steps leading up to and following 7ɑ-dehydration of CA (BaiE) (41), resulting in formation of a 7ɑ-deoxy-3-oxo-Δ^4,6^-intermediates of DCA or LCA, respectively, which are reduced by the BaiH yielding 3-oxo-Δ^4^-intermediates (42). The 3-oxo-Δ^4^-intermediate is then predicted to yield either 3-oxo-DCA (BaiCD) or 3-oxo-allo-DCA (BaiP). The same enzymatic steps are involved in the conversion of CDCA to 3-oxo-Δ^4^-LCA followed by conversion to 3-oxo-LCA (3-oxo-5β-cholan-24-oic acid) or 3-oxo-alloLCA (3-oxo-5ɑ-cholan-24-oic acid) by BaiCD or BaiP, respectively (**Fig. 1**).

We therefore chemically synthesized 3-oxo-Δ^4^-DCA and 3-oxo-Δ^4^-LCA and incubated these substrates (50 μM) with *E. coli* expressing HDCHBGLK_03451 under anaerobic conditions in PBS. When 3-oxo-Δ^4^-LCA was present as the substrate, 3-oxo-allo-LCA (RT = 2.30 min; *m/z* = 373.3) was synthesized, but not 3-oxo-LCA (RT = 2.50 min; *m/z* = 373.3) (**Fig. 2d**). The 6 h reaction yielded 7.00 ± 0.46 μM 3-oxo-allo-LCA (**Fig. 2e**). Similarly, incubation of resting cells with 3-oxo-Δ^4^-DCA yielded a product (RT=1.08 min; *m/z* = 389.26) consistent with 3-oxo-allo-DCA (RT=1.08 min; *m/z* =389.26), but not 3-oxo-DCA (RT= 1.20 min; *m/z* =389.26) (**Fig. 2d**). These data confirm that HDCHBGLK_03451 encodes a novel bile acid 5ɑ-reductase, and we propose the name *baiP* for this gene (See ***SI Appendix*, Fig. S1**).

**Fig. 2.**
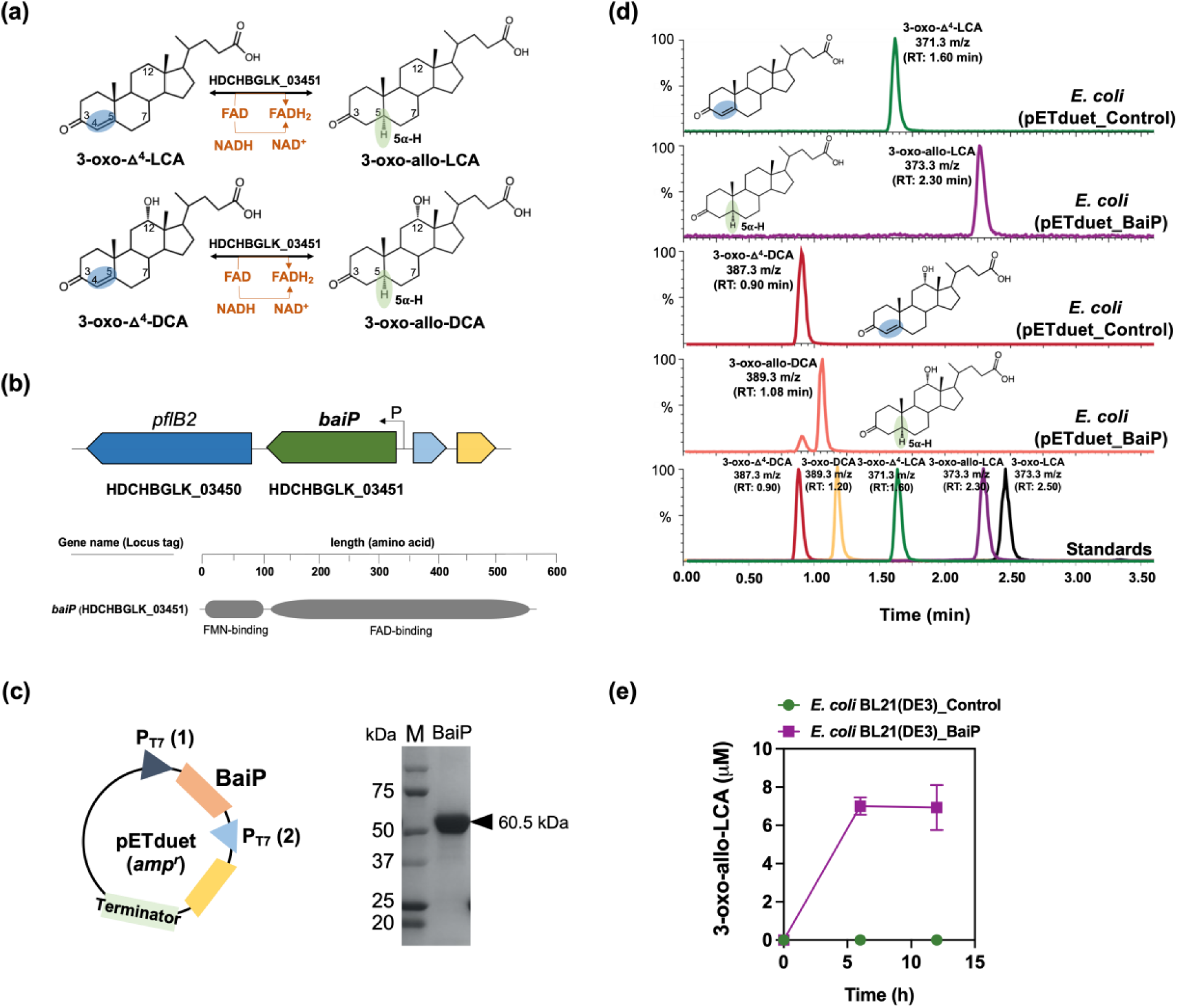
The *baiP* gene from *L. scindens* ATCC 35704 encodes a bile acid 5α-reductase. (a) Formation of bile acid stereoisomers after reduction of 3-oxo-Δ^4^-LCA and 3-oxo-Δ^4^-DCA by 5α-reductase. (b) Gene organization of *baiP* with genomic context and domain structure of BaiP. (c) Cloning strategy for heterologous expression of N-terminal his-tagged recombinant BaiP in *E. coli* BL21(DE3). SDS-PAGE confirms expression of 60.5 kDa recombinant BaiP. (d) Representative LC/MS chromatograms after resting cell assay with *E. coli* BL21(DE3) pETduet_Control or pETduet_BaiP incubated in anaerobic PBS containing 50 μM 3-oxo-Δ^4^-LCA (Top panels 1 & 2) or 50 μM 3-oxo-Δ^4^-DCA (Bottom panels 3 & 4). Standards are shown in Panel 5 (bottom). (e) Time course of 3-oxo-allo-LCA production by the *E. coli* BL21(DE3) pETduet_BaiP strain. Data points indicate the mean concentration of 3-oxo-allo-LCA ± SD (three biological replicates).

We previously reported a cortisol-inducible operon (*desABCD*) in *L. scindens* ATCC 35704 encoding steroid-17,20-desmolase (DesAB) and NADH-dependent steroid 20α-hydroxysteroid dehydrogenase (DesC) (43). DesC reversibly forms cortisol and 20α-dihydrocortisol (43), and DesAB catalyzes the side-chain cleavage of cortisol yielding 11β-hydroxyandrostenedione (11β-OHAD) (43). Because substrates and products in the desmolase pathway have 3-oxo-Δ^4^-structures analogous to 3-oxo-Δ^4^-DCA and 3-oxo-Δ^4^-LCA, we next performed resting cell assays with *E. coli* strain expressing the BaiP enzyme. LC/MS analysis of reaction products indicates that cortisol and 11β-OHAD were not substrates for BaiP (***SI Appendix*, Fig. S2**).

### Phylogenetic analysis of BaiP followed by functional assay reveals the *baiJ* gene also encodes bile acid 5ɑ-reductase

Having provided experimental evidence that *baiP* encodes an enzyme with bile acid 5ɑ-reductase activity, we wanted to determine the phylogeny of the BaiP from *L. scindens* ATCC 35704. A subtree of the >1,400 sequences representing close relatives of the BaiP from *L. scindens* ATCC 35704 was generated (**Fig. 3a**). The proteins most closely related to BaiP from *L. scindens* ATCC 35704 in the “BaiP Cluster’’ were from *Lachnoclostridum* strains MSK.5.24, GGCC_0168, and Lachnospiraceae bacterium 5_1_57FAA. Additional FAD-dependent oxidoreductase BaiP candidates from a penguin isolate, *Proteocatella sphenisci* DSM 23131 (76% sequence identity), and *P. hiranonis* (15, 44) (72% sequence identity) were also identified at high bootstrap values (90-100%). Previous work established *bai* genes in *P. hiranonis* (45), although the present data provide first indication that *P. hiranonis* has the potential to form secondary allo-bile acids (**Fig. 3a, 3b**). *Proteocatella sphenisci* has also been reported to encode the *bai* polycistronic operon (22, 46), and our demonstration that *P. sphenisci* harbors *baiP* indicate that secondary allo-bile acids may constitute part of the bile acid metabolome of penguin guano (**Fig. 3b**).

A second closest FAD-dependent oxidoreductase cluster (∼45% ID) to BaiP from *L. scindens* ATCC 35704 was composed of the previously named BaiJ proteins from *L. scindens* VPI 12708, *L. hylemonae*, and *P. hiranonis*, as well as *Dorea* sp. D27, and an unclassified *Clostridium* sp. (“BaiJ Cluster’’). Previously a novel *bai* operon was identified in which the *baiJ* gene is clustered with *baiK* on a polycistronic operon in *L. scindens* VPI 12708 and *L. hylemonae* DSM 15053 (40). Evidence was also presented that *L. scindens* VPI 12708 and *L. hylemonae* TN271 formed allo-DCA (40). It was then reported that the BaiK is a paralog of BaiF in *L. scindens* VPI 12708, and both proteins catalyze bile acid coenzyme A transferase from the end-product secondary bile acids, DCA∼SCoA and allo-DCA∼SCoA, to primary bile acids including CA, CDCA, allo-CA, and UDCA (23). The *baiJ* gene has been shown previously to be enriched in the gut microbiome in mouse models of liver cancer and CRC (9, 25), diseases reported to be enriched in secondary allo-bile acids in the biliary pool in the few studies that have measured them (47). Taken together, the close phylogenetic clustering of BaiJ with BaiP indicates that the *baiJ* gene may also encode a bile acid 5ɑ-reductase isoform (**Fig. 3a, 3b**) (22, 44, 45).

**Fig. 3.**
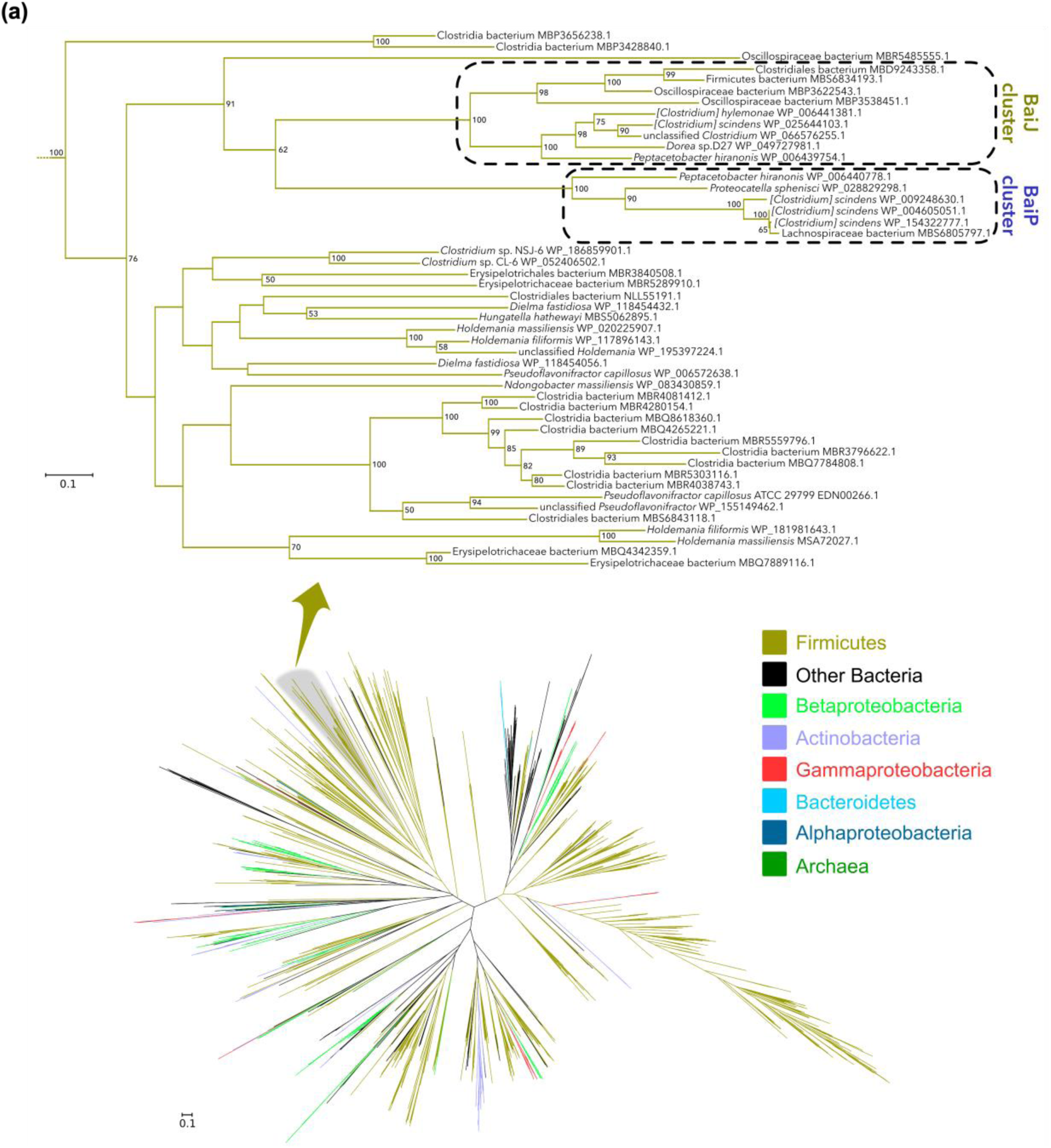
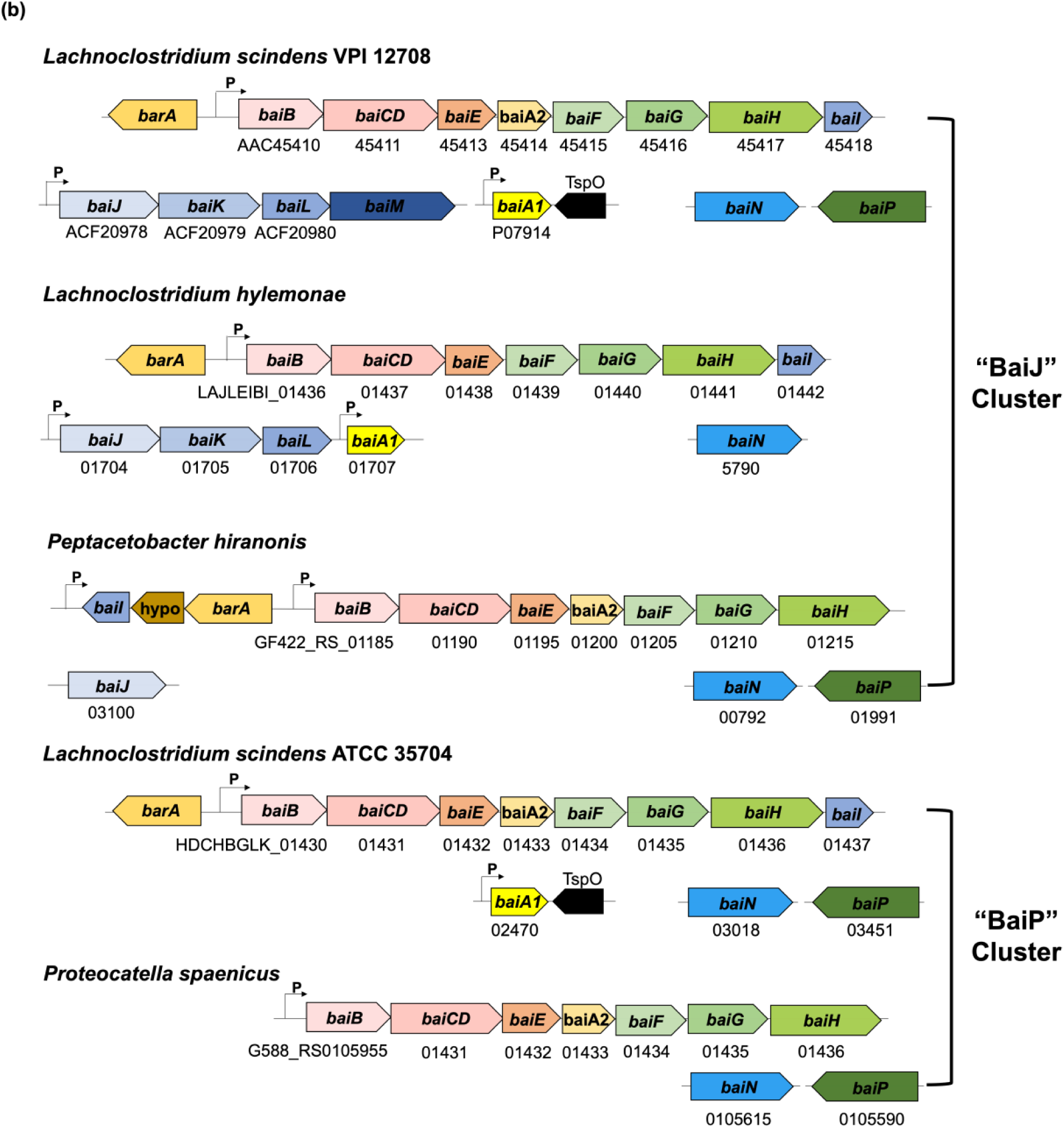
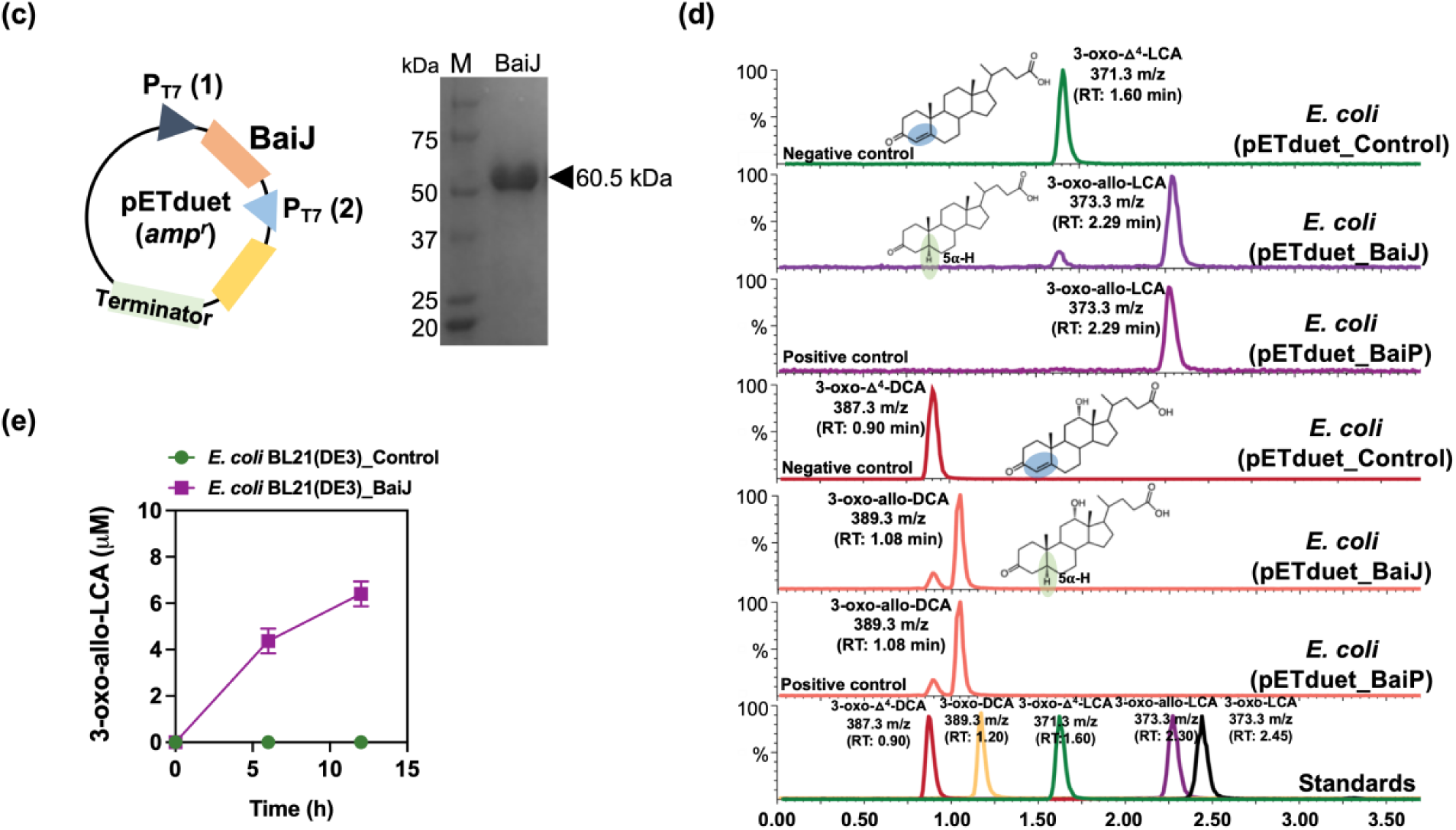
Large scale phylogenetic analysis of BaiP from *L. scindens* ATCC 35704 reveals *baiJ* gene from *C. scindens* VPI 12708 encodes a bile acid 5α-reductase. (a) Maximum-likelihood tree of >2,300 protein sequences from NCBI’s non-redundant database that were similar to BaiP from *L. scindens*. The subtree containing BaiP from *L. scindens* formed two clusters containing BaiP sequences (Purple) from other Firmicutes known to convert CA to DCA. The second cluster contains BaiJ proteins, representing several strains known to convert CA to DCA. (b) Arrangement of genes in the bile acid inducible (*bai*) operon in various species of bile acid 7α-dehydroxylating gut bacteria. The gene encoding enzymes carrying out bile acid metabolism in gut bacteria capable of producing secondary allo-bile acids. Biochemical pathway leading to secondary allo-bile acid formation is shown in Fig. 1. (c) Cloning strategy for *baiJ* gene from *L. scindens* VPI 12708 and SDS-PAGE after purification of recombinant His-tagged BaiJ. (d) Representative LC/MS chromatographs after resting cell assay with E. coli BL21(DE3) pETduet_Control or pETduet_BaiJ incubated in anaerobic PBS containing 50 μM 3-oxo-Δ^4^-LCA (Top panels 1 & 2) compared to pETduet_BaiP (Panel 3). Panels 4 & 5 display chromatograms of reaction products formed after incubation of *E. coli* BL21(DE3) pETduet_Control or pETduet_BaiJ incubated in anaerobic PBS containing 50 μM 3-oxo-Δ^4^-DCA compared to pETduet_BaiP (Panel 6). Standards are shown in Panel 7 (bottom). (e) Time course of 3-oxo-allo-LCA production by the *E. coli* BL21(DE3) pETduet_BaiJ strain. Data points indicate the mean concentration of 3-oxo-allo-LCA ± SD (two biological replicates).

To test this hypothesis, we cloned and overexpressed the *baiJ* gene from *L. scindens* VPI 12708 (accession number: ACF20978) in *E. coli* BL21(DE3) (**Fig. 3c**), and measured conversion of 3-oxo-Δ^4^-LCA and 3-oxo-Δ^4^-DCA in resting cell assays (**Fig. 3d**). When 3-oxo-Δ^4^-LCA (RT = 1.60; *m/z* = 371.25) was the substrate, a product eluting at the same position as 3-oxo-allo-LCA (RT = 2.29; *m/z* = 373.27), but not as 3-oxo-LCA (RT = 2.45; *m/z* = 373.26), was observed. An anaerobic resting cell assay (6 h) resulted in the formation of 4.4 ± 0.54 μM 3-oxo-allo-LCA (**Fig. 3e**). Similarly, when 3-oxo-Δ^4^-DCA (RT = 0.90; *m/z* = 387.25) was the substrate, a product that eluted at the same position as 3-oxo-allo-DCA (RT = 1.08; *m/z* = 389.26), and different from 3-oxo-DCA (RT = 1.20; *m/z* = 389.27), was observed (**Fig. 3d**). These results establish a function for the *baiJ* gene product and indicate that strains of *L. scindens* and other bile acid 7ɑ-dehydroxylating bacteria encode distinct bile acid 5ɑ-reductase isoforms.

### BaiP and BaiA1 catalyze consecutive final reductive steps in the formation of allo-DCA and allo-LCA

Having established that BaiP converts 3-oxo-Δ^4^-LCA to 3-oxo-allo-LCA, we next sought to identify an enzyme from *L. scindens* ATCC 35704 catalyzing the final reductive step from 3-oxo-allo-LCA to allo-LCA. There is compelling evidence that BaiA1 and BaiA2 enzymes catalyze the first oxidative and last reductive steps in the pathway (35). This comes from substrate-specificity and kinetic analyses of BaiA1 and BaiA2 showing that 3-oxo-DCA and 3-oxo-LCA are substrates (48) and by the observation that BaiA is sufficient for the final reductive step yielding DCA (42). Prior work established that the *baiA* genes encode bile acid 3ɑ-hydroxysteroid dehydrogenase (3ɑ-HSDH) that catalyze the first oxidation step, formation of 3-oxo-7ɑ-hydroxy-5β-bile acids, and the final reductive step generating 7-deoxy-3ɑ-hydroxy-5β-bile acids (35). However, the ability of BaiA enzymes to recognize allo-bile acids has not been established (**Fig. 4a**). The *baiA1* gene from *L. scindens* ATCC 35704 was codon-optimized for *E. coli* and overexpressed in *E. coli* alone or in combination with *baiP* (**Fig. 4b**). Whole cell *E. coli* assays with overexpressed BaiA1 converted 3-oxo-allo-LCA (RT = 2.30 min; *m/z* = 373.2) to a product consistent with allo-LCA (RT = 2.74 min; *m/z* = 375.3), but not LCA (RT = 2.68 min; *m/z* = 375.3). *E. coli* expressing both BaiP and BaiA1 converted 3-oxo-Δ^4^-LCA (RT = 1.65 min; *m/z* = 371.3) to allo-LCA (RT = 2.74 min; *m/z* = 375.3) and 3-oxo-Δ^4^-DCA (RT =0.75 min; *m/z* =387.3) to allo-DCA (RT =1.51 min; *m/z* =391.3) confirming the role of BaiP and BaiA1 in the cooperative catalysis of the two final steps in formation of secondary allo-bile acids (**Fig. 4c**).

A previous bioinformatics study claimed based on gene context and annotation that *CLOSCI_00522*, a gene directly downstream from *baiN* (*CLOSCI_00523*), encodes a predicted NAD(FAD)-utilizing dehydrogenase involved in the final reductive step (32) (***SI Appendix*, Fig. S1**). This gene was named “*baiO”* (32). An organism may encode several proteins from different lineages that have similar catalytic activity. Indeed, the BaiN (49) is predicted to catalyze similar sequential reactions to BaiH and BaiCD (42). We therefore tested the hypothesis that the previously annotated *“baiO”* encodes either a bile acid 3-oxo-Δ^4^-reductase and/or bile acid 3ɑ-HSDH. We cloned the *“baiO”* in pETduet and verified the expression after His-tag purification and SDS-PAGE (***SI Appendix*, Fig. S1a, S1b**). Analysis of bile acid products after 24 h incubation of *E. coli* expressing “BaiO” enzyme in a resting cell assay with either 3-oxo-LCA, 3-oxo-DCA (***SI Appendix*, Fig. S1c, S1d)**, 3-oxo-Δ^4^-LCA, or 3-oxo-Δ^4^-DCA (***SI Appendix*, Fig. S1e, S1f)**, did not yield a detectable product by LC/MS. While this does not disprove that *CLOSCI_00522* is involved in bile acid metabolism, we were not able to confirm its function.

**Fig. 4.**
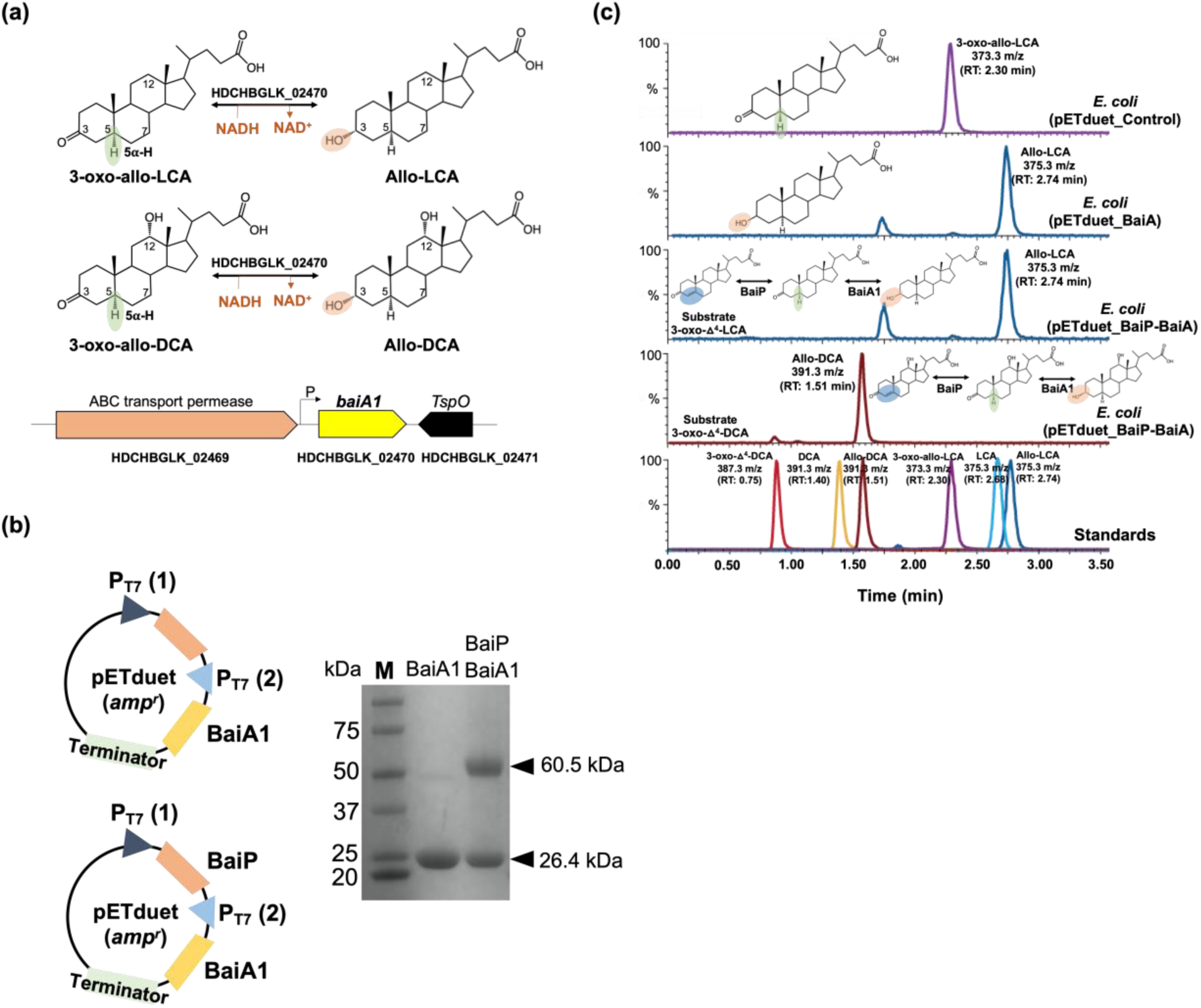
Recombinant BaiA1 from *L. scindens* ATCC 35704 catalyzes the final reductive step in the formation of allo-DCA and allo-LCA. (a) Formation of bile acid stereoisomers after reduction of 3-oxo-allo-LCA and 3-oxo-allo-DCA by 3α-HSDH and gene organization of *baiA1* in *L. scindens* ATCC 35704. (b) Cloning stategy of *baiA1* and *baiA1* + *baiP* in pETduet. SDS-PAGE of His-tagged purified recombinant BaiA1 and BaiA1 + BaiP expressed in *E. coli* BL21(DE3). (c) Representative LC/MS chromatograms after resting cell assay with *E. coli* BL21(DE3) pETduet_Control or pETduet_BaiA1 incubated in anaerobic PBS containing 50 μM 3-oxo-allo-LCA (Top panels 1 & 2), *E. coli* BL21(DE3) pETduet_BaiP-BaiA1 incubated with 50 μM 3-oxo-Δ^4^-LCA (Panel 3) and *E. coli* BL21(DE3) pETduet_BaiP-BaiA1 incubated with 50 μM 3-oxo-Δ^4^-DCA (Panel 4). Standards are shown in Panel 5 (bottom). The overall two-step reaction is shown on the panels.

### The distribution of *baiP* and *baiJ* genes in public human metagenome datasets

Having shown that BaiP clusters with the previously identified BaiJ from *L. hylemonae* DSM 15053, the next objective was to determine the presence of *bai* genes involved in bile acid 7-dehydroxylation among bacterial genomes from human stool samples. We utilized reference sequences of BaiP and BaiJ as well as BaiE and BaiCD (**Fig. 5a**) to generate HMMs in order to search public human metagenomic databases. We expected that the occurrence of BaiE and BaiCD which are co - transcribed on the multi-gene *bai* operon will coincide with the relative abundances of BaiP and BaiJ. As expected, genes for BaiE and BaiCD as well as BaiP and BaiJ were observed to have similar relative frequency (1% and 0.9% of total metagenome assembled genomes (MAGs), respectively). All genes were largely represented by unclassified Firmicutes and *Lachnospiraceae*. (**Fig. 5a)**. Representative genera were analyzed to identify candidates which possess multiple genes of the Bai operon which revealed that unclassified Firmicutes, unclassified *Lachnospiraceae*, and *Flavonifractor* harbored all four genes analyzed. This pathway analysis also revealed the novel finding that *Flavonifractor* and *Pseudoflavonifractor* harbor genes for CA 7-dehydroxylation. Intriguingly, while *bai* genes represented approximately 1% of total MAGs, genes were detected in approximately one third of subjects (BaiCD 35%, BaiE 35%, BaiJ 30%, and BaiP 28%). An analysis of differences in gene presence among healthy subjects and those with adenoma and carcinoma revealed that the genes had the greatest abundance in patients with carcinoma, and that the genes *baiCD, baiE*, and *baiJ* were significantly associated with carcinoma (**Fig. 5b, *SI Appendix*, Table S4, S5**)

## DISCUSSION

The results of the current study add to a growing literature demonstrating that the colonic microbes are capable of “resetting” stereochemistry of sterols undergoing enterohepatic circulation through expression of 5ɑ-reductase and 5β-reductase enzymes. So far, two mechanisms have been identified: (1) A direct mechanism whereby bacteria encoding the multi-step bile acid 7ɑ-dehydroxylation pathway convert primary bile acids to either secondary bile acids via BaiCD/BaiN or as shown herein secondary allo-bile acids via BaiP/BaiJ activities; and (2) an indirect mechanism in which certain species of Bacteroidetes convert 5β-secondary bile acids DCA and LCA to 3-oxo-Δ^4^-intermediates, followed by reduction to secondary allo-bile acids (38). The current work is thus a significant advance towards determining the enzymatic basis for the formation of secondary allo-bile acids by the gut microbiome (**Fig. 6**).

**Fig. 5.**
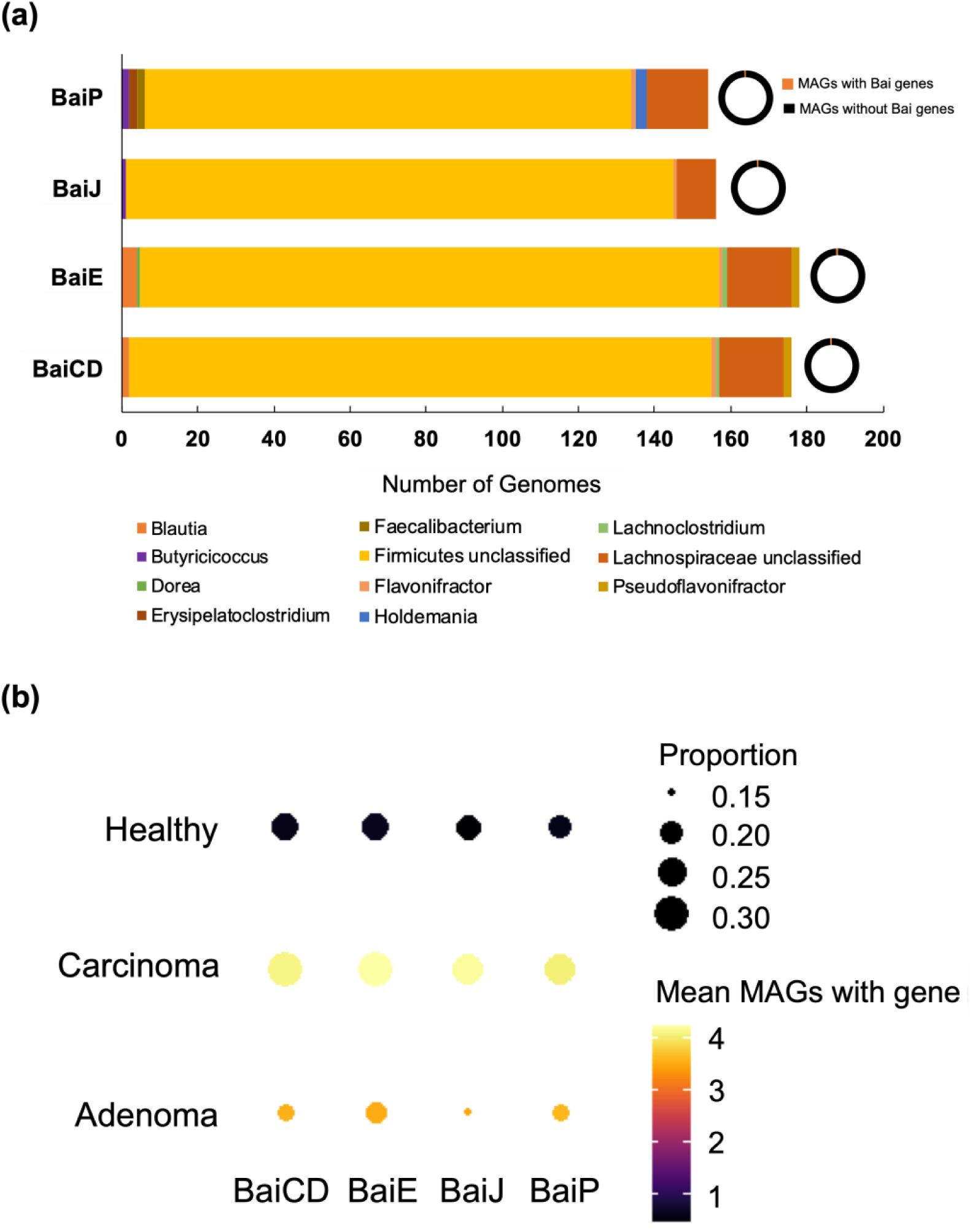
Hidden-Markov Model search reveals enrichment of *bai* genes in colorectal carcinoma. (a) Distribution of microbial genomes with putative 5α-reductase genes (*baiP* and *baiJ*) present across the five metagenomic studies. (b) Dot plots of selected genes related to allo-bile acids production across three disease states: carcinoma, adenoma, and healthy. The size of each dot indicates the proportion of participants with at least one copy of the gene in their bacterial metagenomic assembled genomes (MAGs) and the color of each dot indicates the mean number of MAGs with that gene in the subset of participants that have at least one copy of the gene.

**Fig. 6.**
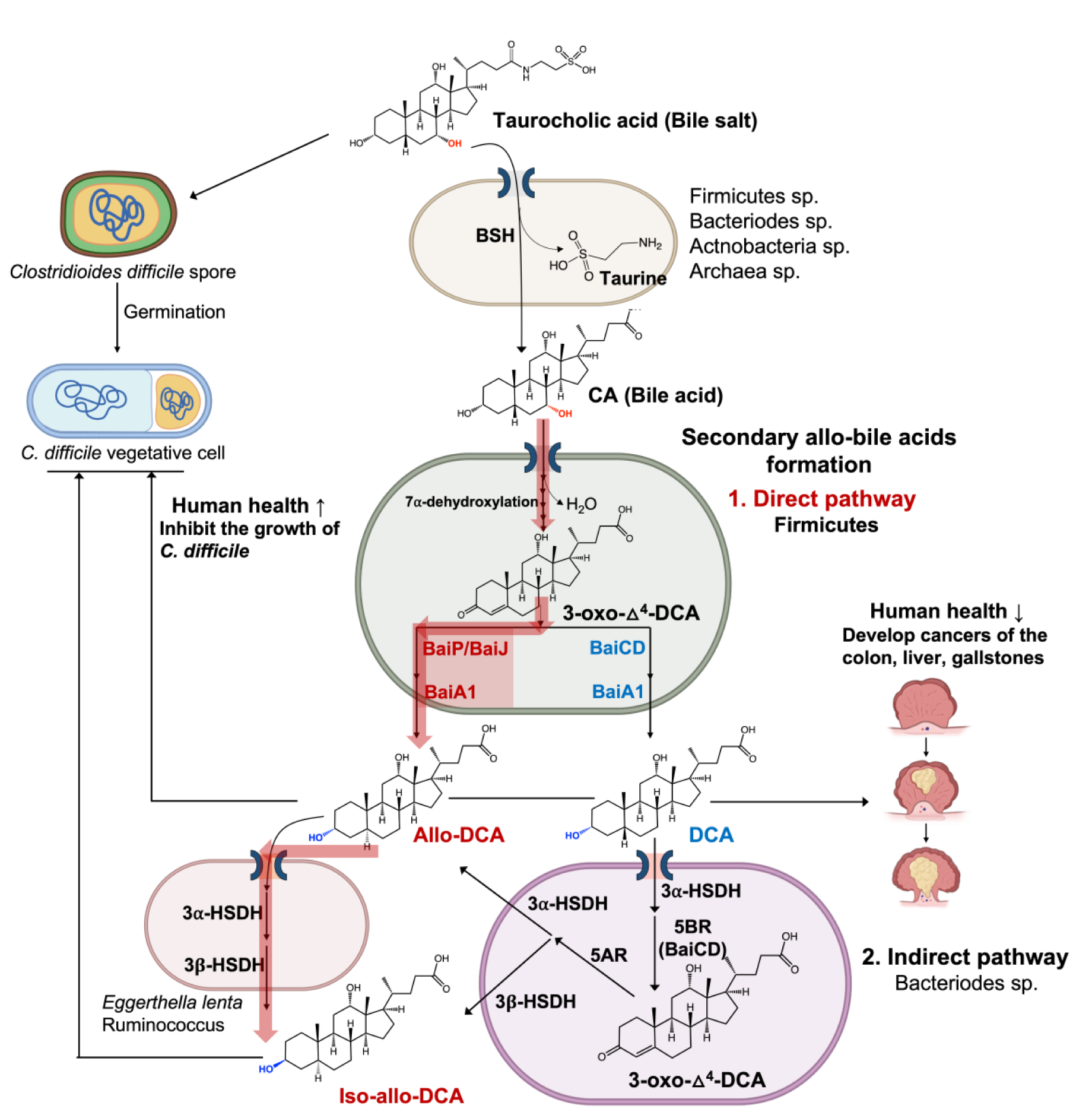
Direct and indirect formation of secondary allo-bile acids, and their potential consequences. Taurocholic acid is deconjugated, mainly in the large intestine, by diverse gut microbial taxa. Free cholic acid is imported into a few species of Firmicutes that harbor the *bai* regulon. **Direct Pathway**: After several oxidative steps, and rate-limiting 7α-dehydration, 3-oxo-4-DCA becomes a substrate for BaiCD forming DCA or BaiP/BaiJ forming alloDCA. **Indirect Pathway**: DCA is imported into Bacteroidetes strains that express 3α-HSDH and 5β-reductase (5BR) which converts DCA to 3-oxo-4-DCA. Expression of 5α-reductase (5AR) and 3beta-HSDH sequentially reduce 3-oxo-4-DCA to iso-alloDCA. Alternatively, alloDCA generated by Firmicutes can be isomerized to iso-alloDCA by species expressing 3α-HSDH and 3β-HSDH such as *Eggerthella lenta*. While taurocholic acid is a germination factor for *C. difficile, s*econdary bile acids such as DCA and secondary allo-bile acids are inhibitory towards *C. difficile* vegetative cells in the GI tract. Secondary bile acids, including DCA and alloDCA, are associated with increased risk of cancers of the liver and colon.

Bile acid intermediates in the 7ɑ-dehydroxylation pathway have been determined previously. Björkhem et al. (20) utilized [3β-^3^H] [24-^14^C] and [5β-^3^H] [24-^14^C] labeled cholic acid in whole cells and cell extracts of *L. scindens* VPI 12708, observing loss of both 3β- and 5β-hydrogens during conversion of CA to DCA (34). Administration of [3β-^3^H] [24-^14^C] CA to volunteers followed by analysis of tritium loss after extraction from duodenal aspirates confirmed that 3-oxo-Δ^4^-bile acid intermediates were formed during conversion of CA to DCA (34). Subsequent work incubating [24-^14^C] CA with cell extracts of *L. scindens* VPI 12708 revealed a multi-enzyme pathway necessary to convert CA to DCA (and CDCA to LCA) (50). Hylemon and Bjӧrkhem (1991) isolated nine [24-^14^C] CA intermediates after incubation with cell-free extracts of CA-induced whole cells of *L. scindens* VPI 12708 providing the biochemical framework to search for enzymes involved in bile acid 7ɑ-dehydroxylation (34). Subsequent work determined that bile acid 7ɑ-dehydroxylation proceeds by two oxidation steps yielding a 7ɑ-hydroxy-3-oxo-Δ^4^-intermediate, the substrate for the rate-limiting enzyme, bile acid 7ɑ-dehydratase (BaiE) (35, 41). Removal of the C7-hydroxyl yields a 7-deoxy-3-oxo-Δ^4,6^-intermediate which is then reduced by flavoproteins BaiN (49) or BaiH (42) to a 7-deoxy-3-oxo-Δ^4^-intermediate. The BaiCD and BaiA isoforms then convert 7-deoxy-3-oxo-Δ^4^-intermediates to DCA or LCA (35, 42, 51) One of the bile acid-inducible [24-^14^C] CA metabolites identified was [24-^14^C] allo-DCA, indicating that *L. scindens* possesses an enzyme with bile acid 5ɑ-reductase distinct from BaiCD (bile acid 5β-reductase) (34). The current results establish conclusively that the *baiP* and *baiJ* genes encode bile acid 5ɑ-reductases in different strains of *L. scindens* and related Firmicutes that catalyze the formation of allo-DCA and allo-LCA.

Previous work also demonstrated that BaiA1 and BaiA2 catalyze both the initial oxidation and final reduction in the formation of DCA and LCA (41,42). However, a recent report named a gene (*CLOSCI_00522*) adjacent to *baiN*, the “*baiO*’’ that encodes a predicted 61 kDa flavin-dependent dehydrogenase proposed to catalyze the final reductive step in the pathway (32). We tested both BaiA1 and BaiO for reduction of allo-DCA and allo-LCA. While the function of *CLOSCI_00522* in bile acid metabolism remains unclear, our results have extended the functional role of the BaiA1. We determined for the first time that this enzyme converts 3-oxo-allo-DCA and 3-oxo-allo-LCA to allo-DCA and allo-LCA, respectively.

The functional role of the previously reported *baiJKL* operon in *L. scindens* VPI 12708 and *L. hylemonae* TN271 has also been extended by the current study (23). Ridlon and Hylemon (2012) reported that BaiK and BaiF catalyze bile acid∼CoA transferase from secondary bile acids, including allodeoxycholyl∼SCoA, to primary bile acids (23). The *baiJ* gene was annotated as “flavin-dependent fumarate reductase” and “3-ketosteroid-Δ^1^-dehydrogenase”, and is co-expressed with *baiKL* under the control of the conserved *bai* promoter (23). We previously observed bile acid induction of *baiJKL* genes by RT-PCR (50) and RNA-Seq (52) in *L. hylemonae* TN271. Also, the *baiJ* gene was reported to be enriched in the gut microbiome in mouse models of liver cancer and CRC (9,25). Fecal secondary allo-bile acids have also been reported to be enriched in GI cancers (47).

Phylogenetic analysis of BaiP from *L. scindens* ATCC 35704 revealed two clusters harboring Firmicutes encoding the *bai* pathway, many of which, such as *P. hiranonis, L. hylemonae*, and strains of *L. scindens*, are known to convert CA and CDCA to DCA and LCA, respectively. These clusters are also represented by taxa such as *Dorea* sp. D27, *P. sphenisci*, and *Oscillospiraceae* MAGs whose genome sequences contain *bai* operons (45, 46). Clusters with more distant homologs of BaiP are also worth examining in future studies for novel bile acid 3-oxo-Δ^4^-reductases. Mining human metagenomic datasets for “core” Bai proteins (BaiCD, BaiE) as well as BaiP and BaiJ sequences confirmed that these enzymes are only encoded in Firmicutes. Roughly a third of healthy, adenoma, and carcinoma subjects had detectable BaiE enzymes representing ∼1% of MAGs. A combination of low abundance bile acid 7-dehydroxylating Firmicutes and stringency of the HMM search likely explains the low representation of subjects with detectable Bai enzymes. Intriguingly, and in line with previous reports^25^, Bai enzymes are enriched in CRC subjects relative to healthy subjects.

There is a paucity of studies on secondary allo-bile acids, and the literature which exists is conflicting as to whether to regard these hydrophobic “flat” bile acids as beneficial, disease promoting, or contextually important (47). Recent work measured the secondary allo-bile acid iso-allo-LCA in fecal samples at an average concentration of ∼20 μM, and that low micromolar levels, such as those achieved in our resting cell assays, inhibit the growth of gram-positive pathogens including *C. difficile* (38) (**Fig. 6**). There is a recent growing interest in the immune mechanisms of action of secondary bile acid derivatives and isomers in the colon. Secondary bile acid derivatives, including 3-oxo-DCA, 3-oxo-LCA, iso-DCA (3β, 12ɑ-dihydroxy-5β-cholan-24-oic acid), iso-LCA (3β-hydroxy-5β-cholan-24-oic acid), and certain secondary allo-bile acids (e.g. iso-allo-LCA: 3β-hydroxy-5ɑ-cholan-24-oic acid), regulate the balance of regulatory T cells (Treg) and pro-inflammatory T_H_17 cells by promoting expansion of Tregs (53-55). The current work is thus an important contribution in a rapidly evolving area of the role of diverse bile acid metabolites generated by the gut microbiome on mechanisms underlying host health and disease.

## MATERIALS AND METHODS

### Bacterial strains and chemicals

*E. coli* Top10 [F-*mcr*A Δ(*mrr*-*hsd*RMS-*mcr*BC) φ80*lac*ZΔM15 Δ*lac*X74 *rec*A1 *ara*D139 Δ(*ara*-*leu*) 7697 *gal*U *gal*K *rps*L (Str^R^) *end*A1 *nup*G] competent cells from Invitrogen (Carlsbad, CA, USA) were used for manipulation of plasmids, and *E. coli* BL21(DE3) [F−, *ompT, hsdSB*(rB− mB−), *gal, dcm, rne131* (DE3)] was also purchased from Invitrogen and used for protein expression. 3-oxo-Δ^4^-LCA, 3-oxo-allo-LCA, 3-oxo-LCA, allo-LCA, LCA, and 3-oxo-DCA were purchased from Steraloids (Newport, RI, USA). Isopropyl β-D-1-thiogalactopyranoside (IPTG) was purchased from Gold Biotechnology (St. Louis, MO, USA). All other reagents were of the highest possible purity and purchased from Fisher Scientific (Pittsburgh, PA, USA).

### Bile acid synthesis

Authentic 3-oxo-Δ^4^-DCA and allo-DCA were synthesized as previously described (56) and confirmed by nuclear magnetic resonance (NMR) spectroscopy (***SI Appendix*, Fig. S3, S4**).

### Cloning of *bai* operon genes from *L. scindens* strains

The strains/plasmids, primers, and synthetic DNA sequences used in this study are listed in ***SI Appendix*, Table S1, S2**, and **S3**, respectively. First, *baiP* gene encoding FAD-dependent oxidoreductase and *baiA1* gene encoding 3α-HSDH from *L. scindens* ATCC 35704, *baiJ* gene encoding FAD-dependent oxidoreductase from *L. scindens* VPI 12708, and “*baiO*” encoding a predicted 61 kDa flavin-dependent dehydrogenase were codon-optimized for *E. coli* and synthesized using gBlocks service from Integrated DNA Technologies (IDT, IA, USA). To construct a BaiP, BaiJ, “BaiO” or BaiA1 expression plasmid (pBaiP, pBaiJ, pBaiO or pBaiA1), a DNA fragment (vector fraction) was amplified from the pETduet plasmid using a primer pair of V1-F and V1-R, V1-F and V1-R, V1-F and V1-R, or V2-F and V2-R, respectively. Another DNA fragment (insert fraction) was amplified from the synthetic oligomers of BaiP, BaiJ, “BaiO” or BaiA1 using a primer pair of BaiP-F and BaiP-R, BaiJ-F and BaiJ-R, “BaiO”-F and “BaiO”-R or BaiA1-F and BaiA1-R, respectively. The two pairs of PCR products were ligated together by *in vitro* homologous recombination using a Gibson assembly cloning kit (NEB, Boston, MA, USA), respectively. For construction of a BaiP and BaiA1 co-expression plasmid (pBaiP-A1), a DNA fragment (vector fraction) was amplified from the pBaiP plasmid using a pair of the primers V2-F and V2-R, and another DNA fragment (insert fraction) was amplified from the synthetic oligomer of BaiA1 using a pair of the primers BaiA1-F and BaiA1-R. The two PCR products were ligated together by the Gibson assembly cloning kit (NEB)

Recombinant plasmids (***SI Appendix*, Table S1**) were transformed into chemically competent *E. coli* Top10 cells via heat-shock method, respectively, plated, and grown for overnight at 37° on lysogeny broth (LB) agar plates supplemented with appropriate antibiotics (Ampicillin: 100 μg/ml). A single colony from each transformation was inoculated into LB medium (5 ml) containing the corresponding antibiotic. The cells were subsequently centrifuged (3,220×g, 10 min, 4°) and plasmids were extracted from the cell pallets using QIAprep Spin Miniprep kit (Qiagen, CA, USA). The sequences of the inserts were confirmed by Sanger sequencing (ACGT Inc, Wheeling, IL, USA).

### Heterologous expression and purification of Bai enzymes in *E. coli*

For protein expression, the extracted recombinant plasmids were transformed into *E. coli* BL21(DE3) cells by use of electroporation method, respectively, and cultured overnight at 37° on LB agar plates supplementary with appropriate antibiotics. Selected colonies were inoculated into 10 mL of LB medium containing the corresponding antibiotic and grown at 37° for 6 h with vigorous aeration. The pre-cultures were added to fresh LB medium (1L), supplemented with appropriate antibiotics, and aerated at 37° until reaching an OD_600_ (optical density of a sample measured at a wavelength of 600 nm) of 0.3. IPTG was added to each culture at a final concentration of 0.1 mM to induce and the temperature was decreased to 16°. Following 16 h of culturing, cells were pelleted by centrifugation (4000×g, 30 min, 4°) and resuspended in 30 ml of binding buffer (20 mM Tris-HCl, 300 mM NaCl, 10 mM 2-mercaptoethanol, pH 7.9). The cell suspension was subjected to an ultra sonicator (Fisher Scientific) and the cell debris was separated by centrifugation (20,000×g, 40 min, 4°).

The recombinant protein in the soluble fraction was then purified using TALON Metal Affinity Resin (Clontech Laboratories, CA, USA) per manufacturer’s protocol. The recombinant protein was eluted using an elution buffer composed of 20 mM Tris-HCl, 300 mM NaCl, 10 mM 2-mercaptoethanol, and 250 mM imidazole at pH 7.9. The resulting purified protein was analyzed using sodium dodecyl sulfate-polyacrylamide gel electrophoresis (SDS-PAGE).

### Whole cell bile acid conversion assay

*E. coli* BL21(DE3) strains harboring the constructed plasmids were cultured aerobically at 25° on LB medium (10 mL) supplementary with appropriate antibiotics and expressed the corresponding proteins by IPTG induction at 25°. Following 16 h of culturing, the strains were pelleted by centrifugation (3,220×g, 10 min) and washed twice with anaerobic PBS solution. The washed *E. coli* strains were inoculated along with 50 μM bile acid substrates (3-oxo-Δ^4^-LCA, 3-oxo-Δ^4^-DCA, or 3-oxo-allo-LCA) into 10 mL of PBS and incubated anaerobically at room temperature for 12 h. The whole cell reaction cultures were centrifuged at 3,220×g for 10 min to remove bacterial cells and adjusted the pH of the supernatant to pH 3.0 by adding 25 μL of 2N HCl. Bile acid metabolites were extracted by vortexing with two volumes of ethyl acetate for 1 to 2 min. The organic layer was recovered and evaporated under nitrogen gas. The products were dissolved in 200 μL methanol and analyzed by liquid chromatography-mass spectrometry (LC-MS).

### Liquid chromatography-mass spectrometry

LC-MS analysis for all samples was performed using a Waters Acquity UPLC system coupled to a Waters SYNAPT G2-Si ESI mass spectrometer (Milford, MA, USA). For the bile acids as substrates and products of whole cell bioconversion assay by the *E. coli* strains expressing BaiP, BaiJ, or BaiP-A1 enzymes (3-oxo-Δ^4^-LCA, 3-oxo-Δ^4^-DCA, 3-oxo-LCA, 3-oxo-allo-LCA, 3-oxo-DCA, LCA, allo-LCA, DCA, and allo-DCA) analysis, LC was performed with a Waters Acquity UPLC HSS T3 C18 column (1.8 μm particle size, 2.1 mm × 100 mm) at a column temperature of 40°. Samples were injected at 0.2 μL. Mobile phase A was a mixture of acetonitrile and methanol (50/50, v/v), and B was 10mM ammonium acetate. The mobile phase composition was 75% of mobile phase A and 25% of mobile phase B and ran an isocratic mode. The flow rate of the mobile phase was 0.5 mL/min. MS was carried out in negative ion mode with a desolvation temperature of 400° and desolvation gas flow of 800 L/hr. The capillary voltage was 2,000 V. Source temperature was 120°, and the cone voltage was 30V. Chromatographs and mass spectrometry data were analyzed using Waters MassLynx software. Analytes were identified according to their mass and retention time. For quantification of 3-oxo-allo-LCA produced by the *E. coli* BL21(DE3) expressing BaiP/BaiJ strains, a standard curve was obtained, and then 3-oxo-allo-LCA was quantified based on the standard curve (**Fig. S5**). The limit of detection (LOD) for 3-oxo-Δ^4^-LCA, 3-oxo-allo-LCA, and allo-LCA was 0.1 μmol/L.

For the cortisol and 11β-OHAD as substrates and products of whole cell bioconversion assay by the *E. coli* strain expressing BaiP enzyme analysis, LC was performed with a Waters Acquity UPLC BEH C18 column (1.7 μm particle size, 2.1 mm × 50 mm) at a column temperature of 40°. Samples were injected at 0.2 μL. Mobile phase A was a mixture of 95% water, 5% acetonitrile, and 0.1% formic acid, and B was a mixture of 95% acetonitrile, 5% water, and 0.1% formic acid. The mobile phase gradient was as follows: 0 min 100% mobile phase A, 0.5 min 100% A, 6.0 min 30% A, 7.0 min 0% A, 8.1 min 100% A, and 10.0 min 100% A. The flow rate of the mobile phase was 0.5 mL/min. MS was carried out in positive ion mode with a desolvation temperature of 450° and desolvation gas flow of 800 L/hr. The capillary voltage was 3,000 V. Source temperature was 120°, and the cone voltage was 30V.

### NMR spectroscopy

To determine the molecular structure of the chemically synthesized 3-oxo-Δ^4^-DCA and allo-DCA at the atomic level, NMR spectroscopy was performed. ^1^H-NMR spectra were recorded on a JNM-ECA-500 spectrometer (JEOL Co., Tokyo, Japan) at 500 MHz, with pyridine-D_5_ as the solvent. Chemical shifts are given as the δ-value with tetramethylsilane (TMS) as an internal standard. The abbreviation used here: s, singlet; d, doublet; bs, broad singlet.

### Phylogenetic Analysis

Sequences for phylogenetic analyses were retrieved from NCBI’s NR protein database using the sequence of HDCHBGLK_03451 as the query and limiting the number of resulting database matches to five thousand and allowing a maximum alignment E-value of 1E-10 for BLASTP v. 2.12.0+ (57). The retrieved alignments showed high sequence conservation, therefore the worst E-value seen in the alignments was about 3E-37.

Given the high sequence similarities observed in the search step, sequences were clustered with USEARCH v. 11.0.667 (58) to remove redundancy from the dataset. The cluster_fast command was used with an identity threshold of at least 95% to cluster sequences. Each cluster was represented in the phylogenetic analysis by one representative, the centroid sequence. The only exception was the sequences in the same cluster as the query sequence used above, in which case all sequences from the cluster were used in the analysis, instead of just the centroid. Clustering resulted in 1,603 sequences included in the downstream analyses.

Centroids 25% shorter or longer than the average sequence length calculated for the whole dataset (596 amino acids) were removed from the dataset, thus keeping in the analysis only sequences with at least 446 and at most 744 amino acids in length. The 1,460 protein sequences remaining in the dataset were aligned by MUSCLE v. 3.8.1551 (59) and the best-fitting sequence substitution model was identified using ModelTest-NG v. 0.1.7 (60). Phylogenetic tree inference was performed using the maximum likelihood criterion as implemented by RAxML v. 8.2.12 (61), using the WAG sequence substitution model with empirical residue frequencies, gamma-distributed substitution rates, and bootstrap pseudoreplicates (whose number, 250, was determined automatically by the program at run-time). The resulting phylogenetic tree was edited with TreeGraph2 v. 2.15.0-887 (62) and Dendroscope v. 3.7.6 (63) and further cosmetic adjustments were performed with the Inkscape vector editor (https://inkscape.org/ last accessed on January, 20th, 2022).

### Bai gene identification in MAG database

A database of publicly available MAGs from five cohorts varying in CRC status was previously annotated for open reading frames and used for this study (64,65). Custom Hidden Markov Model (HMM) profiles were created for each of the 4 genes of interest (*baiCD, baiE, baiP*, and *baiJ*) by creating an alignment of reference protein sequences in this study and blastp results with 60% identity to those reference sequences and then passing the alignments to hmmbuild to create an HMM profile. Initial HMM cutoffs were generated by querying protein sequences from the Human Microbiome Project (64). To further refine HMM profile cutoffs, blast databases were made of each alignment and a concatenated file of predicted open reading frames from the 16,936 MAGs described earlier were queried against the alignment databases. The MAG database was searched using the HMM profiles with finalized cutoffs and hmmsearch within HMMER 3.1b2. All custom HMM profiles used for these searches can be found at: https://github.com/escowley/BileAcid_LeeJ.

### Summary calculations and statistical analysis for association of Bai genes with disease state from MAG database

Summary calculations of number of gene hits in the MAG database, number of participants with the gene of interest, and disease information were performed in R and can be found in ***SI***

***Appendix*, Table S4**. Methods for determining associations between Bai genes and disease state were previously described (63). Briefly, chi squared tests were performed on a dataset of binarized participants that were designated as “presence” if any of their MAGs contained a copy of the gene of interest or “absence” if none of their recovered MAGs contained a copy of the gene of interest. P-values less than 0.05 are designated as significant (***SI Appendix*, Table S5**).

## Acknowledgements

We would like to acknowledge financial support from National Institutes of Health grants (1RO1 CA204808-01 [J.M.R, H.R.G.], R01 GM134423-01A1 [J.M.R], R03 AI147127-01A1 [J.M.P.A, J.M.R]) as well as UIUC Department of Animal Sciences Matchstick grant, and Hatch ILLU-538-916. ESC is a Medical Scientist Training Program (MSTP) student and was supported by a National Library of Medicine training grant to the Computation and Informatics in Biology and Medicine Training Program (NLM 5T15LM007359) at UW-Madison, and in part by MSTP grant T32 GM140935. PGW was supported at UI-Chicago by the Cancer Education and Career Development Program grant T32CA057699. LKL was supported by NSF GRFP 2017224867. HLD was supported by the David H. and Norraine A. Baker Graduate Fellowship in Animal Sciences at Illinois.

## Notes

### Competing Interest Statement

The authors have declared no competing interest.

## References

[1] F. Hofmann, The continuing importance of bile acids in liver and intestinal disease. Arch. Intern. 159, 2647–2658 (1999).

[2] T. Midtvedt, Microbial bile acid transformation. Am J Clin Nutr 27, 1341–1347 (1974).

[3] J. R. Swann et al., Systemic gut microbial modulation of bile acid metabolism in host tissue compartments. Proc. Natl. Acad. Sci.108, 4523–4530 (2011).

[4] S. Narushima et al., Deoxycholic acid formation in gnotobiotic mice associated with human intestinal bacteria. Lipids 41, 835–843 (2006).

[5] B. E. Gustafsson, T. Midtvedt, A. Norman, Metabolism of cholic acid in germfree animals after the establishment in the intestinal tract of deconjugating and 7 alpha-dehydroxylating bacteria. Acta Pathol. Microbiol. Scand. 72, 433–443 (1968).

[6] K. Lan et al., Key role for the 12-hydroxy group in the negative ion fragmentation of unconjugated C24 bile acids. Anal. Chem. 88, 7041–7048 (2016).

[7] R. A. Quinn et al., Global chemical effects of the microbiome include new bile-acid conjugations. Nature 579, 123–129 (2020).

[8] H Zhou, P. B. Hylemon, Bile acids are nutrient signaling hormones. Steroids 86, 62–68 (2014).

[9] S. Yoshimoto et al., Obesity-induced gut microbial metabolite promotes liver cancer through senescence secretome. Nature 499, 97–101 (2013).

[10] C. Ma et al., Gut microbiome-mediated bile acid metabolism regulates liver cancer via NKT cells. Science 360 (2018).

[11] S. Ocvirk, S. J. O’Keefe, Influence of bile acids on colorectal cancer risk: Potential mechanisms mediated by diet-gut microbiota interactions. Curr. Nutr. Rep. 6, 315–322 (2017).

[12] S. N. Chaudhari et al., Bariatric surgery reveals a gut-restricted TGR5 agonist with antidiabetic effects. Nat. Chem. Biol.17, 20–29 (2021).

[13] J. Y. L. Chiang, J. M. Ferrell, Discovery of farnesoid X receptor and its role in bile acid metabolism. Mol. Cell Endocrinol. 548, 111618 (2022).

[14] F. Berr, G. A. Kullak-Ublick, G. Paumgartner, W. Münzing, P. B. Hylemon, 7 alphadehydroxylating bacteria enhance deoxycholic acid input and cholesterol saturation of bile in patients with gallstones. Gastroenterology 111, 1611–1620 (1996).

[15] J. E. Wells, F. Berr, L. A. Thomas, R. H. Dowling, P. B. Hylemon, Isolation and characterization of cholic acid 7alpha-dehydroxylating fecal bacteria from cholesterol gallstone patients. J. Hepatol. 32, 4–10 (2000).

[16] J. Marksteiner, I. Blasko, G. Kemmler, T. Koal, C. Humpel, Bile acid quantification of 20 plasma metabolites identifies lithocholic acid as a putative biomarker in Alzheimer’s disease. Metabolomics 14, 1 (2018).

[17] MahmoudianDehkordi et al., Altered bile acid profile associates with cognitive impairment in Alzheimer’s disease - An emerging role for gut microbiome. Alzheimers Dement 15, 76–92 (2019).

[18] G. Porez, J. Prawitt, B. Gross, B. Staels, Bile acid receptors as targets for the treatment of dyslipidemia and cardiovascular disease. J. Lipid Res. 53, 1723–1737 (2012).

[19] B. V. M. Begley, C. Hill, C. G. Gahan, J. R. Marchesi, Functional and comparative metagenomic analysis of bile salt hydrolase activity in the human gut microbiome. Proc. Natl. Acad. Sci. 105, 13580–13585 (2008).

[20] Björkhem, K. Einarsson, P. Melone, P. Hylemon, Mechanism of intestinal formation of deoxycholic acid from cholic acid in humans: Evidence for a 3-oxo-delta 4-steroid intermediate. J. Lipid Res. 30, 1033–1039 (1989).

[21] J. M. Ridlon, S. C. Harris, S. Bhowmik, D.-J. Kang, P. B. Hylemon, Consequences of bile salt biotransformations by intestinal bacteria. Gut Microbes 7, 22–39 (2016).

[22] K. H. Kim et al., Identification and characterization of major bile acid 7α-dehydroxylating bacteria in the human gut. mSystems 10.1128/msystems.00455-22, e0045522 (2022).

[23] J. M. Ridlon, P. B. Hylemon, Identification and characterization of two bile acid coenzyme A transferases from Clostridium scindens, a bile acid 7α-dehydroxylating intestinal bacterium. J. Lipid Res. 53, 66–76 (2012).

[24] V. L. Hale et al., Shifts in the fecal microbiota associated with adenomatous polyps. Cancer Epidemiol. Biomarkers Prev. 26, 85–94 (2017).

[25] J. Wirbel et al., Meta-analysis of fecal metagenomes reveals global microbial signatures that are specific for colorectal cancer. Nat. Med. 25, 679–689 (2019).

[26] T. Ochsenkühn et al., Colonic mucosal proliferation is related to serum deoxycholic acid levels. Cancer 85, 1664–1669 (1999).

[27] A van Faassen et al., Plasma deoxycholic acid is related to deoxycholic acid in faecal water. Cancer Lett. 114, 293–294 (1997).

[28] E. Bayerdörffer et al., Unconjugated secondary bile acids in the serum of patients with colorectal adenomas. Gut 36, 268–273 (1995).

[29] E. Bayerdörffer et al., Increased serum deoxycholic acid levels in men with colorectal adenomas. Gastroenterology 104, 145–151 (1993).

[30] J. Ou et al., Diet, microbiota, and microbial metabolites in colon cancer risk in rural Africans and African Americans. Am. J. Clin. Nutr. 98, 111–120 (2013).

[31] S. Ocvirk et al., A prospective cohort analysis of gut microbial co-metabolism in Alaska Native and rural African people at high and low risk of colorectal cancer. Am. J. Clin. Nutr.111, 406–419 (2020).

[32] A Heinken et al., Systematic assessment of secondary bile acid metabolism in gut microbes reveals distinct metabolic capabilities in inflammatory bowel disease. Microbiome 7, 75 (2019).

[33] C Kwan, I. Frouni, D. Bédard, A. Hamadjida, P. Huot, Granisetron, a selective 5-HT3 antagonist, reduces L-3,4-dihydroxyphenylalanine-induced abnormal involuntary movements in the 6-hydroxydopamine-lesioned rat. Behav. Pharmacol. 32, 43–53 (2021).

[34] P. B. Hylemon, P. D. Melone, C. V. Franklund, E. Lund, I. Björkhem, Mechanism of intestinal 7 alpha-dehydroxylation of cholic acid: Evidence that allo-deoxycholic acid is an inducible sideproduct. J. Lipid Res. 32, 89–96 (1991).

[35] S. Bhowmik et al., Structure and functional characterization of a bile acid 7α dehydratase BaiE in secondary bile acid synthesis. Proteins: Struct. Funct. Genet. 84, 316–331 (2016).

[36] T. Tadano, M. Kanoh, M. Matsumoto, K. Sakamoto, T. Kamano, Studies of serum and feces bile acids determination by gas chromatography-mass spectrometry. Rinsho Byori 54, 103–110 (2006).

[37] T. Tadano et al., Kinetic analysis of bile acids in the feces of colorectal cancer patients by gas chromatography-mass spectrometry (GC-MS). Rinsho Byori 55, 417–427 (2007).

[38] Y. Sato et al., Novel bile acid biosynthetic pathways are enriched in the microbiome of centenarians. Nature 599, 458–464 (2021).

[39] S. Devendran et al., Clostridium scindens ATCC 35704: Integration of nutritional requirements, the complete genome sequence, and global transcriptional responses to bile acids. Appl. Environ. Microbiol. 85, e00052–00019 (2019).

[40] J. M. Ridlon, D.-J. Kang, P. B. Hylemon, Isolation and characterization of a bile acid inducible 7α-dehydroxylating operon in Clostridium hylemonae TN271. Anaerobe 16, 137–146 (2010).

[41] J. A. Dawson, D. H. Mallonee, I. Björkhem, P. B. Hylemon, Expression and characterization of a C24 bile acid 7 alpha-dehydratase from Eubacterium sp. strain VPI 12708 in Escherichia coli. J. Lipid Res. 37, 1258–1267 (1996).

[42] M. Funabashi et al., A metabolic pathway for bile acid dehydroxylation by the gut microbiome. Nature 582, 566–570 (2020).

[43] J. M. Ridlon et al., Clostridium scindens: A human gut microbe with a high potential to convert glucocorticoids into androgens. J. Lipid Res. 54, 2437–2449 (2013).

[44] X. J. Chen et al., Characterization of Peptacetobacter hominis gen. nov., sp. nov., isolated from human faeces, and proposal for the reclassification of Clostridium hiranonis within the genus Peptacetobacter. Int. J. Syst. Evol. Microbiol. 70, 2988–2997 (2020).

[45] J. E. Wells, P. B. Hylemon, Identification and characterization of a bile acid 7αdehydroxylation operon in Clostridium sp. strain TO-931, a highly active 7α-dehydroxylating strain isolated from human feces. Appl. Environ. Microbiol. 66, 1107–1113 (2000).

[46] M. Vital, T. Rud, S. Rath, D. H. Pieper, D. Schlüter, Diversity of bacteria exhibiting bile acidinducible 7α-dehydroxylation genes in the human gut. Comput. Struct. Biotechnol. J. 17, 1016–1019 (2019).

[47] S. J. Shiffka, M. A. Kane, P. W. Swaan, Planar bile acids in health and disease. Planar bile acids in health and disease. Biochim Biophys Acta Biomembr 1859, 2269–2276 (2017).

[48] S. Bhowmik et al., Structural and functional characterization of BaiA, an enzyme involved in secondary bile acid synthesis in human gut microbe. Proteins 82, 216–229 (2014).

[49] S. C. Harris et al., Identification of a gene encoding a flavoprotein involved in bile acid metabolism by the human gut bacterium Clostridium scindens ATCC 35704. Biochim. Biophys. Acta. Mol. Cell Biol. Lipids 1863, 276–283 (2018).

[50] J. M. Ridlon, D.-J. Kang, P. B. Hylemon, Bile salt biotransformations by human intestinal bacteria. J. Lipid Res. 47, 241–259 (2006).

[51] J. Kang, J. M. Ridlon, D. R. Moore, 2nd, S. Barnes, P. B. Hylemon, Clostridium scindens baiCD and baiH genes encode stereo-specific 7alpha/7beta-hydroxy-3-oxo-delta4-cholenoic acid oxidoreductases. Biochim. Biophys. Acta. 1781, 16–25 (2008).

[52] J. M. Ridlon et al., The ‘in vivo lifestyle’ of bile acid 7α-dehydroxylating bacteria: comparative genomics, metatranscriptomic, and bile acid metabolomics analysis of a defined microbial community in gnotobiotic mice. Gut Microbes 11, 381–404 (2020).

[53] X. Song et al., Microbial bile acid metabolites modulate gut RORγ^+^ regulatory T cell homeostasis. Nature 577, 410–415 (2020).

[54] W. Li et al., A bacterial bile acid metabolite modulates Treg activity through the nuclear hormone receptor NR4A1. Cell Host Microbe 29, 1366-1377.e1369 (2021).

[55] S. Hang et al., Bile acid metabolites control T_H_17 and T_reg_ cell differentiation. Nature 576, 143–148 (2019).

[56] R. A. Leppik, Improved synthesis of 3-keto, 4-ene-3-keto, and 4,6-diene-3-keto bile acids. Steroids 41, 475–484 (1983).

[57] Camacho et al., BLAST+: Architecture and applications. BMC Bioinform. 10, 421 (2009).

[58] R. C. Edgar, Search and clustering orders of magnitude faster than BLAST. Bioinformatics 26, 2460–2461 (2010).

[59] R. C. Edgar, MUSCLE: A multiple sequence alignment method with reduced time and space complexity. BMC Bioinform. 5, 113 (2004).

[60] D Darriba et al., ModelTest-NG: A new and scalable tool for the selection of DNA and protein evolutionary models. Mol. Biol. Evol. 37, 291–294 (2019).

[61] A Stamatakis, RAxML version 8: A tool for phylogenetic analysis and post-analysis of large phylogenies. Bioinformatics 30, 1312–1313 (2014).

[62] C. Stöver, K. F. Müller, TreeGraph 2: Combining and visualizing evidence from different phylogenetic analyses. BMC Bioinform. 11, 7 (2010).

[63] H. Huson, C. Scornavacca, Dendroscope 3: An interactive tool for rooted phylogenetic trees and networks. Syst. Biol. 61, 1061–1067 (2012).

[64] P. G. Wolf et al., Diversity and distribution of sulfur metabolic genes in the human gut microbiome and their association with colorectal cancer. Microbiome 10, 64 (2022).

[65] E. Pasolli et al., Extensive unexplored human microbiome diversity revealed by over 150,000 genomes from metagenomes spanning age, geography, and lifestyle. Cell 176, 649-662.e620 (2019).

